# A web portal for gene expression across all life stages of *Schistosoma mansoni*

**DOI:** 10.1101/308213

**Authors:** Z. Lu, Y. Zhang, M. Berriman

## Abstract

RNA-seq approach can provide useful information about gene expression. Although several studies have been conducted in the parasite *Schistosoma mansoni*, the gene expression data is often limited to differential analysis between certain life stages. A recent meta-analysis of RNA-seq studies generated valuable expression data across all life stages of *S. mansoni*. To facilitate the use and visualisation of these data, we established an interactive web portal implementing not only data from above-mentioned analysis, but also functional aspects including conserved domains and associated pathways, as a complement to main databases for *S. mansoni*. Users can also visualise and analyse their own data via the web portal. The interactive visualisation implemented in the web portal can facilitate characterising schistosome genes for the research community.

## Introduction

Schistosomes affect almost 240 million people worldwide (WHO). There is no vaccine, and only one commonly-used drug against these parasites. As drug resistance may arise (Doenhoff et al. 2008), it is important to find novel drug targets, for which gene expression profiling can be very useful (Chengalvala et al. 2007). RNA-seq is among the most common tools for determining gene expression levels, either absolute or relative. For schistosomes, various RNA-seq studies have been performed in the past, focusing on certain life stages. Currently, main databases for schistosomes, including GeneDB, Wormbase Parasite or SchistoDB, have not implemented or only partial of those expression data.

A recent meta-analysis of published RNA-seq experiments provides a comprehensive view of gene expression changes across all life stages of *Schistsoma mansoni* (Lu and Berriman 2018). However, as it’s not possible to cover all genes in one publication, it’s very likely that there are more interesting candidates that were not mentioned in the analysis. To facilitate the use of these additional results, we present here a web portal to visualise all expression data from the meta-analysis, as well as functional aspects about the gene, such as protein domains and associated KEGG pathways. Furthermore, users can create their own interactive chart and link their own data to the results from the meta-analysis or to any external databases, which will be able to accelerate analysing their data.

## Methods

### Implementation

As shown in Fig. 1, the web portal integrates different types of data into a typical gene page. The users can generate their own interactive chart and link the data to internal gene pages or external pages.

**Figure 1.**
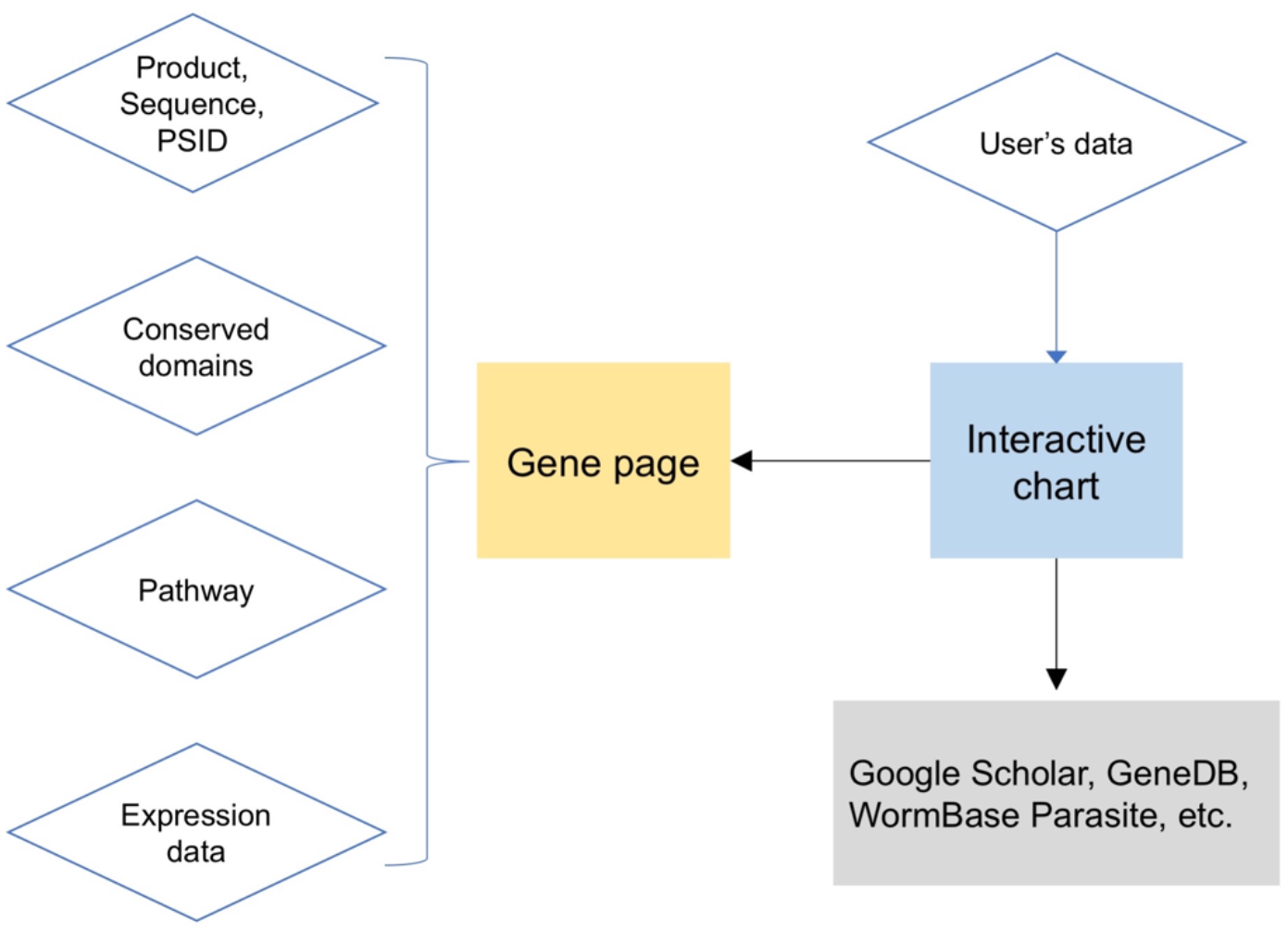
Schematic view of the web portal design. Detailed data acquisition and analysis pipeline were mentioned below.

Expression data was generated based on the meta-analysis of RNA-seq data covering all life stages of *Schistosoma mansoni*. RPKM values calculated after counts normalisation were used to reveal the expression profile of each gene and indicated as “Normalised expression”. Scatter plots and line charts were produced using the open-source chart editor IVIS (Lu and Zhang 2017).

Protein sequences, and information on product and previous identifiers were obtained from GeneDB (http://www.genedb.org/Homepage/Smansoni;accessedon10/07/2017).

Conserved protein domains were analysed using NCBI Batch CD-Search (https://www.ncbi.nlm.nih.gov/Structure/bwrpsb/bwrpsb.cgi; analysed on 08/08/2017; E-Value threshold 0.01, max. hits 500) using the CDD database (Marchler-Bauer et al. 2017). KEGG pathway database (Kanehisa and Goto 2000) using KAAS (http://www.genome.jp/kaas-bin/kaas_main; analysed on 08/08/2017; program: GHOSTZ, alignment method: BBH).

### Operation

The users can query the web portal by the Smp gene id (e.g., Smp_000020) or the product description (e.g., receptor). By either mean the user will be directed to the result page with hyperlinks to individual gene pages.

To create their own charts, users only need to prepare the data in comma-, space-, or tab-separated format. The rows and columns can be switched before charting, if necessary. The links for each data point can be configured to any database pages.

For all charts in the web portal, users can zoom in to see underlying data. The charts can be printed or downloaded into different formats.

### Use Cases

As shown in Fig. 2A, expression data, significance testing for different pairwise comparisons, as well as basic information about the gene, e.g., conserved domains, orthologue and associated pathways, will be available on a typical gene page. Besides, each differential expression analysis will be provided as an interactive chart, where zooming and tooltips indicating gene id, log-transformed fold-change and FDR values are functional. In addition, each gene id is connected with the corresponding gene page (Fig. 2B). Finally, users will be able to create their own expression plot, and to link their own data to the resulting pages from the meta-analysis, or any other databases, for speedy data filtration (Fig. 2C).

**Figure 2.**
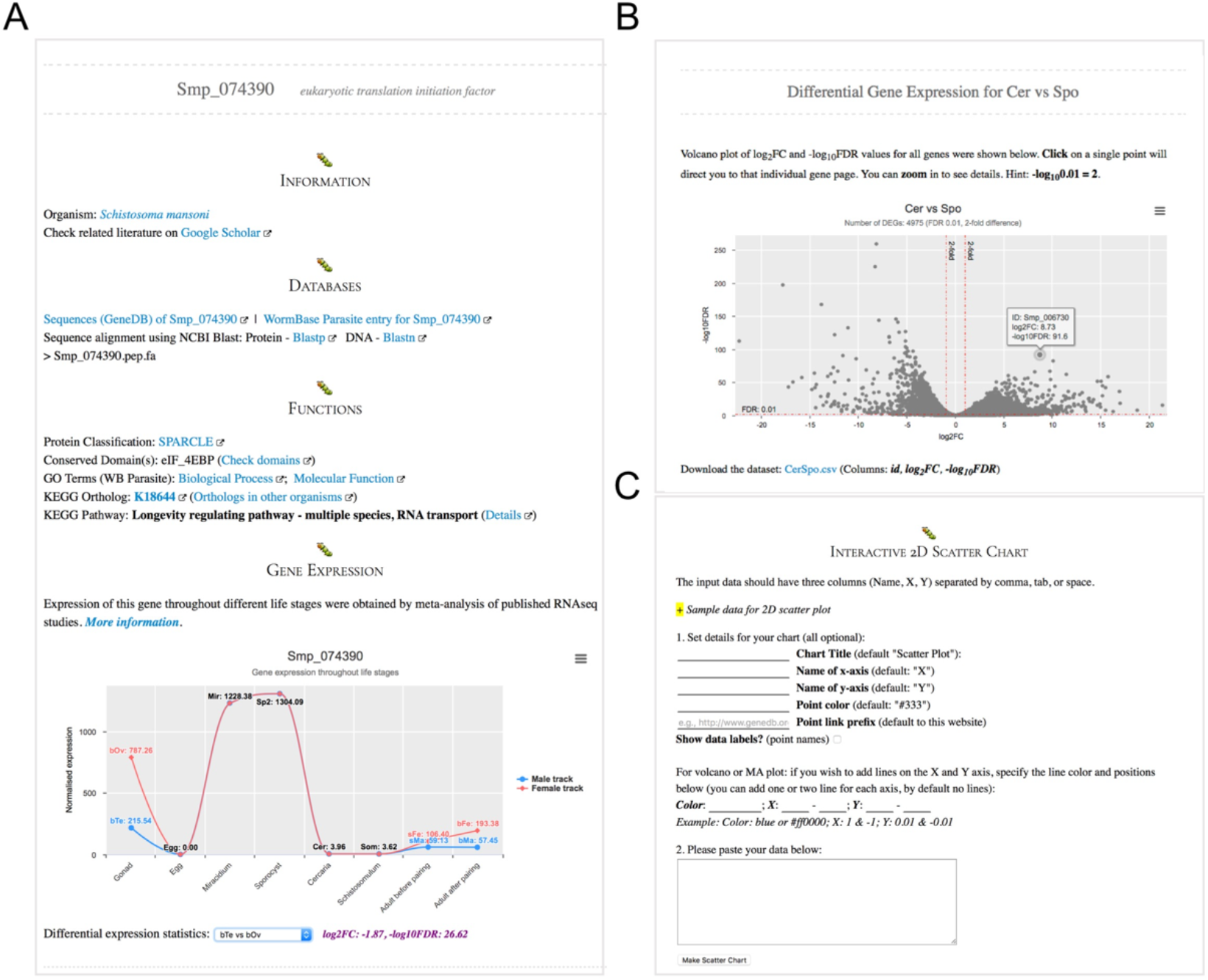
Exemplary pages of the web portal. (A) An exemplary gene page, which includes functional (e.g., conserved domains and pathways) and expression data about the gene. (B) Differential expression data interface. Zooming and hyperlinks are integrated. (C) The charting unit. Line and scatter charts can be produced, and users can link their own data to gene pages or external databases.

## Discussion

We provide there a web portal for accessing gene expression data across all life stages of *S. mansoni*, and in addition a tool to generate interactive charts using the users’ own data. While the web portal is helpful in accessing and analysing gene expression, there are some limitations:

1. As the genome annotation might change after some time, the identifiers in the web portal might be out-dated. This is currently the biggest limitation but can be solved by updating the database regularly, e.g., every three months. Apart from that, the users can always link their own data to the latest database;
2. If the genome assembly version is updated, the data needs to be re-analysed, which will involve some effort. However, this doesn’t mean that all expression data or the tendency will be changed.
3. Although the presented web portal is mainly demonstrating data from the meta-analysis, similar structural framework can be integrated into main databases for schistosomes such as WormBase Parasite, which is a desirable feature in the future development of the pipeline.

## Data and Software Availability

Web portal to access all expression data: https://meta.schisto.xyz

## Competing Interests

The authors declare no competing interests.

